# Modeling Microtubule-Cytoplasm Interaction in Plant Cells

**DOI:** 10.1101/2022.09.29.510138

**Authors:** Mohammad Murshed, Donghui Wei, Ying Gu, Jin Wang

**Affiliations:** Department of Mathematics, University of Tennessee at Chattanooga, Chattanooga, TN 37403, USA; Department of Biochemistry and Molecular Biology, Pennsylvania State University, State College, PA 16801, USA

**Keywords:** microtubule organization, fluid-structure interaction, computational modeling

## Abstract

Although microtubules in plant cells have been extensively studied, the mechanisms that regulate the spatial organization of microtubules are poorly understood. We hypothesize that the interaction between microtubules and cytoplasmic flow plays an important role in the assembly and orientation of microtubules. To test this hypothesis, we developed a new computational modeling framework for microtubules based on theory and methods from the fluid-structure interaction. We employed the immersed boundary method to track the movement of microtubules in cytoplasmic flow. We also incorporated details of the encounter dynamics when two microtubules collide with each other. We verified our computational model through several numerical tests before applying it to the simulation of the microtubule-cytoplasm interaction in a growing plant cell. Our computational investigation demonstrated that microtubules are primarily oriented in the direction orthogonal to the axis of cell elongation. We validated the simulation results through a comparison with the measurement from laboratory experiments. We found that our computational model, with further calibration, was capable of generating microtubule orientation patterns that were qualitatively and quantitatively consistent with the experimental results. The computational model proposed in this study can be naturally extended to many other cellular systems that involve the interaction between microstructures and the intracellular fluid.

## 1 Introduction

Microtubules (MTs) are filamentous intracellular structures that form part of the cytoskeleton within the cytoplasm. They play a significant role in the growth of plant cells [12,42,52]. Each microtubule has a fast-growing plus end with exposed *β*-tubulin and a slow-growing minus end with exposed *α*-tubulin [29]. It exhibits a distinct behavior of dynamic instability by switching between growth and shrinkage during cell expansion [7, 25, 48].

One of the most intriguing properties of microtubules is that they typically form parallel arrays beneath the inner surface of the plasma membrane. It has been found that in an elongating wild-type plant cell, the cortical MT arrays are predominantly arranged orthogonal to the cell elongation axis [66]. Moreover, a well-known ‘direct guidance model’ [37] postulates that microtubules guide the deposition of cellulose microfibrils, the major load-bearing polymers in the cell wall. This hypothesis has been supported by numerous observations that newly synthesized microfibrils mirror the orientation of microtubules [2, 72]. Thus, the overall assembly and orientation of cortical MT arrays appear to be essential in regulating the mechanical properties of the cell wall and in controlling the anisotropic expansion of plant cells [42]. However, the fundamental mechanisms shaping the spatial organization of microtubules into such specific patterns remain poorly understood.

Computational models have long been providing useful insight into biology [50, 62]. Mathematical modeling of plant cell growth started many decades ago when Lockhart developed the first explicit model for cell elongation by connecting turgor pressure and cell wall extensibility [46]. Since then the Lockhart equation, together with many variations, have become a standard modeling paradigm for plant cell growth and cell wall mechanics [24, 41, 53, 56, 59, 63, 65]. There are also a number of modeling and simulation studies specifically devoted to microtubule dynamics [1, 9, 13, 18, 33, 35, 36, 47, 51, 54, 64]. These studies have successfully represented the dynamic instability and treadmilling behavior of a microtubule for its shrinkage at the minus end and growth at the plus end [39,71]. They have also extensively investigated the contact dynamics between two microtubules with three possible collision outcomes: zippering (when the encounter angle < 40°), induced catastrophe and cross-over (when the encounter angle > 40°) [14, 19, 64].

Although these findings have greatly improved our understanding in MT dynamics, they have not provided sufficient insight into the mechanisms that regulate the organization and orientation of microtubules [11, 13, 44, 45, 71]. Particularly, none of these models have considered the interaction between cytoplasm and microtubules. Cytoplasm, enclosed by the plasma membrane, mainly consists of water that takes up over 80% of the cell volume. Microtubules are immersed in cytoplasm, and their motion directly impacts each other. During the period of rapid cell growth, cytoplasm is driven and accelerated by turgor pressure, pushing outward on the membrane and exerting forces on microtubules, which could significantly impact the dynamical behavior of microtubules. We hypothesize that the interaction between microtubules and cytoplasmic flow plays an important role in the spatial organization of microtubules. The main objective of the present work is to test this hypothesis through computational modeling.

Given the ubiquitous presence of fluids in plants, fluid mechanics have been modeled and simulated in plant biology [58]. There have been extensive mathematical modeling studies on fluid flow in plants at macroscales, such as the transpiration stream, water flow across tissues, and xylem and phloem transport [16, 22, 30, 55]. However, within a single plant cell, fluid dynamics have been rarely investigated [10, 65], with very few published results. For example, a fluid model was proposed in [17] to study cell wall development during plant cell growth. Another fluid study in [40] analyzed concentration boundary layers of osmotically driven flows in plant cells. In addition, cytoplasmic streaming in plant cells has been modeled and its role in aiding transport and mixing of molecular contents has been analyzed [27]. To our best knowledge, no study has been performed for the impact of fluid dynamics on the spatial organization of microtubules.

The present paper represents a pilot effort toward filling this knowledge gap. We have developed a new mathematical and computational framework for the cytoplasm-microtubule interaction within a plant cell, utilizing the notion of fluid-structure interaction (FSI) [15] where the cytoplasm is treated as a viscous fluid and each microtubule is treated as a solid structure immersed in the fluid. Our findings, based on a two-dimensional (2D) modeling framework, provide new insight through the angle of fluid dynamics into the spatial organization and orientation of microtubules. This study could enable subsequent model extension to three-dimensional (3D) space.

## 2 Materials and Methods

### 2.1 FSI model

Our computational model for the FSI study of microtubules is based on the immersed boundary method, originally proposed by Peskin to solve the Navier-Stokes equations with moving boundaries [57]. This numerical technique solves the fluid equations with an additional forcing term that represents the effects of the immersed structure on the fluid motion. The fluid equations are computed in a fixed Eulerian mesh, which avoids expensive mesh-updating procedures. The immersed structure is represented as a moving boundary which is tracked on a separate Lagrangian mesh. The immersed boundary method and its many variants have become one of the most popular approaches for FSI computation, due to their efficiency, flexibility and robustness [32, 38, 49].

To facilitate model development and algorithm implementation, we focus on a two-dimensional (2D) setting as an approximation to the real intracellular scenario, and represent each microtubule as an elastic fiber. In fact, it has been observed that the motion of microtubules is mostly confined to a 2D surface of the cytoplasmic face beneath the plasma membrane [13, 29].

We first describe the formulation of a general 2D FSI model that can be later applied to the study of the cytoplasm-microtubule dynamics. Here we are concerned with the interaction between an incompressible viscous fluid and *M* elastic fibers (*M* ≥ 1) immersed in the fluid to be computed by utilizing the immersed boundary method [57, 70]. We consider the fluid in a two-dimensional domain Ω. We assume that each fiber has approximately a zero volume and is represented by a one-dimensional curve (or, boundary) Γ_*m*_ for 1 ≤ *m* ≤ *M*. The motion of the fluid is described by the Navier-Stokes equations [3], the most fundamental equations in fluid dynamics:

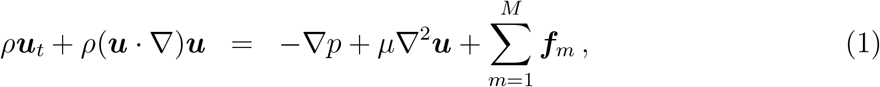

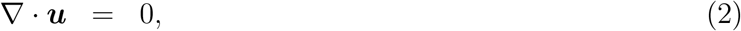

where (1) is the momentum equation and (2) is the continuity equation (also called the incompressibility condition). The variable ***u*** is the velocity vector and *p* is the pressure. The parameters *ρ* and *μ* are the density and the dynamic viscosity, respectively, of the fluid. The last term in the momentum equation, 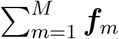, represents the total force that the elastic fibers exert on the flowhich is referred to as the FSI force.

In the immersed boundary framework, the fluid motion is described by the Eulerian approach and equations (1) and (2) are solved on a fixed Cartesian mesh. We let ***x*** = (*x*, *y*) denote the Eulerian coordinate in the fluid domain Ω. On the other hand, the structural motion is tracked by the Lagrangian approach. We let ***X***_*m*_(*s*, *t*) denote the position of the *m*th fiber at time *t*, where *s* is a variable to parameterize the boundary Γ_*m*_. Let also ***F***_*m*_(*s*, *t*) be the force density of the *m*th fiber at time *t*. We then have

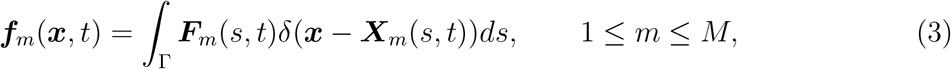

where *δ* denotes the Dirac delta function. Equation (3) shows that the FSI force ***f***_*m*_(***x***, *t*) is singular; i.e., it is zero everywhere in the fluid domain Ω except on the boundary Γ_*m*_.

Each fiber interacts with the viscous fluid and moves with the local fluid at the same velocity due to the no-slip condition. We thus have

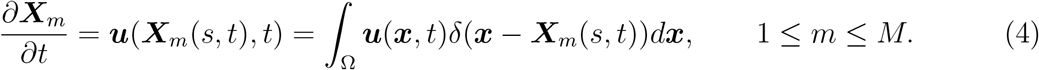

Equations (3) and (4) are used to communicate the force and velocity information between the fluid and the structure.

We denote the numerical solution for the velocity, the pressure, and the FSI force from the *m*th fiber (1 ≤ *m* ≤ *M*) as 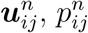, and 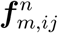, respectively, at the location ***x***_*ij*_ = (*x*_*i*_, *y*_*j*_) and time *t_n_*. Suppose that *K* Lagrangian points (*K* ≥ 2) are used to discretize each curve Γ_*m*_ for 1 ≤ *m* ≤ *M*. We let 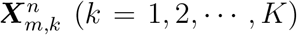 denote the *k*th point on Γ_*m*_ at time *t_n_*, and 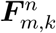 and 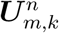 denote the force density and structural velocity, respectively, at the point 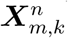.

The two integral equations (3) and (4) can then be written as discrete sums,

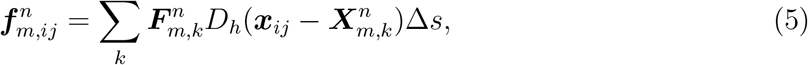

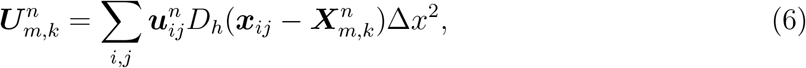

where *D*_*h*_ is a discrete approximation to the 2D delta function with *h* = Δ*x* = Δ*y*. A common choice for the 2D discrete delta function is

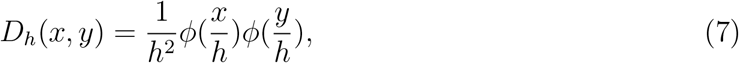

where *ϕ* is the one-dimensional discrete delta function derived in [31, 57]:

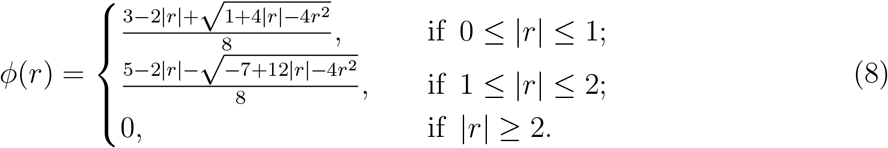

Equation (5) distributes the force from the structure to the fluid mesh, whereas equation (6) interpolates the fluid velocity to the structural points.

### 2.2 Force calculation

A fiber can be represented as a sequence of movable nodes, where the line joining a node to one of its neighboring nodes is called a link. As the fiber is flexible, it is able to undergo stretching deformation as well as bending deformation. Such deformations generate the spring force and the bending force [4, 20, 21].

Consider two consecutive nodes ***X***_*MA*_ = (*x*_*MA*_, *y*_*MA*_) and ***X***_*SL*_ = (*x*_*SL*_, *y*_*SL*_), which may be referred to as the ‘master’ and ‘slave’ nodes, respectively. The spring force between these two nodes is computed as

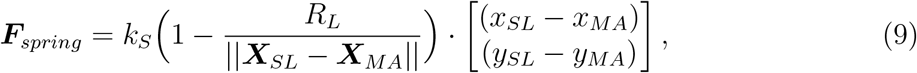

where *k*_*S*_ is the stretching stiffness constant and *R*_*L*_ the resting length between ***X***_*MA*_ and *X*_*SL*_.

Consider three consecutive nodes ***X***_*L*_ = (*x*_*L*_, *y*_*L*_), ***X***_*M*_ = (*x*_*M*_, *y*_*M*_), and ***X***_*R*_ = (*x*_*R*_, *y*_*R*_), which are referred to as the left, middle and right nodes, respectively. The bending force that acts on ***X***_*M*_ is then calculated as

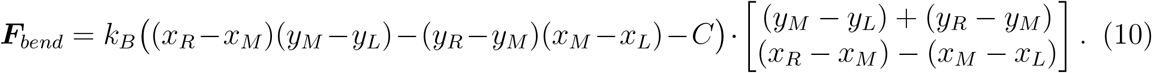

The parameter *k*_*B*_ is the bending stiffness constant, and *C* is the curvature that can be calculated as

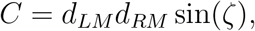

where *d*_*LM*_ is the length of the link between ***X***_*L*_ and ***X***_*M*_, *d*_*RM*_ is the length of the link between ***X***_*M*_ and ***X***_*R*_, and *ζ* is the angle at the relaxed state.

We let ***F***_*add*_ denote any additional force acting on each fiber. For example, if we take the mass of the fibers into account, then the gravitational force

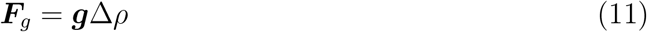

needs to be considered, where Δ*ρ* = *ρ*_*fiber*_ − *ρ*_*fluid*_ and ***g*** is the gravitational acceleration. Then we have ***F***_*add*_ = ***F***_*g*_. In such a case, it is typically assumed that Δ*ρ* ≪ *ρ*_*fluid*_ so that the Boussinesq approximation can be applied; i.e., the average density of the fluid-fiber mixture is approximately the same as that of the fluid [70].

The overall force density on each fiber is then determined by the spring force, the bending force, and the additional force:

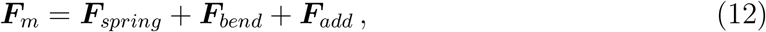

for 1 ≤ *m* ≤ *M*.

### 2.3 Numerical procedure

Now we describe the procedure to implementing the immersed boundary method for the coupled fluid-fiber system. Given the numerical solution at time *t*_*n*_, the following steps advance the solution to time *t*_*n*+1_

1. Use equation (12) to compute the force density on each fiber based on current boundary positions and deformations.
2. Spread the force from the Lagrangian nodes for the fibers to the Eulerian lattice points for the fluid using equation (5).
3. With the FSI force computed from Step 2, solve the Navier-Stokes equations on the Eulerian grid and update the velocity and pressure. This is typically accomplished by using the projection method [8] where an intermediate velocity ***u***\* is first calculated from the momentum equation,

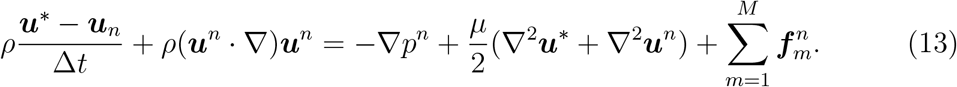

The intermediate velocity is then projected into a divergence-free subspace to compute the new velocity and pressure at time *t_n_*_+1_,

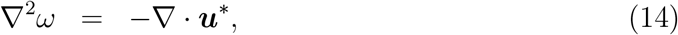

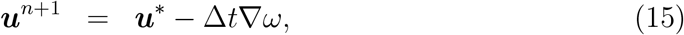

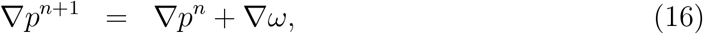

where *ω* is referred to as the pressure increment and computed from the Poisson equation (14).
4. Use equation (6) to compute the structural velocity from the local fluid velocity. Then update the positions of the structural points on each fiber:

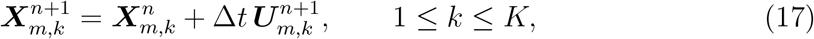

for 1 ≤ *m* ≤ *M*.

### 2.4 Microtubule encounter dynamics

Microtubules constantly encounter and collide with each other during their movement in a plant cell, which directly impacts their assembly and organization. Since we are mainly concerned with microtubules as the fibers in our computational model, we will also incorporate the detail of the MT encounter dynamics in our FSI study. The interaction between microtubules has been routinely investigated through fluorescent images. Based on experimental results from time-lapse fluorescence microscopy, three different modes of collision and self-organization are observed for microtubules [14]:

1. **Zippering** (also called ‘bundling’): A microtubule collides at a shallow angle (< 40°) with a pre-existing microtubule. The encountering microtubule subsequently changes its initial trajectory to align with the impeding microtubule.
2. **Induced catastrophe** (also called ‘collision-induced depolymerization’): A microtubule encounters a pre-existing microtubule at a steep angle (≥ 40°) and then depolymerizes and shortens after the collision.
3. **Cross-over**: An encountering microtubule approaches a pre-existing microtubule at a steep angle (≥ 40°) and subsequently crosses over the impeding microtubule.

Since we are concerned with 2D simulation, the cross-over mode is not applicable. Meanwhile, prior studies have suggested that zippering and induced catastrophe may have a larger contribution to MT alignment and organization than cross-over does [13, 18]. We thus focus our attention on the effects of zippering and induced catastrophe in this work. The two types of behaviors are distinguished by the angle of collision between the encountering MT (EMT) and the pre-existing MT (PEMT). Figure 1 illustrates the collision between the two microtubules, where the angle Φ_2_ can be computed via the law of cosine:

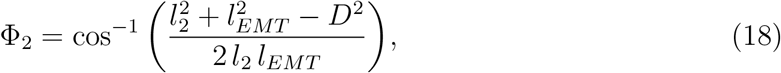

where *l*_*EMT*_ is the length of the EMT, and *D* is the distance between the tail of the EMT and that of the PEMT. If Φ_2_ < 40°, then zippering will take place. If 40° ≤ Φ_2_ ≤ 90°, then catastrophe will take place. Additionally, if Φ_2_ > 90°, then we examine the angle Φ_1_ = 180° − Φ_2_ instead and use the length *l*_1_ instead of *l*_2_ to compute Φ_1_.

**Figure 1:**
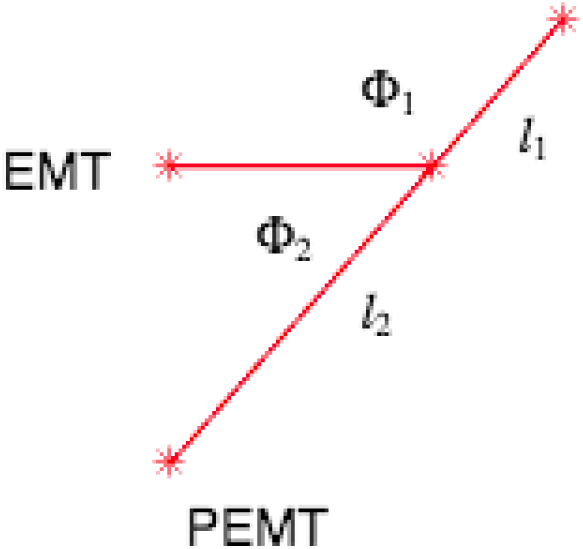
The angle of collision determines the encountering dynamics between two microtubules, where EMT refers to the encountering microtubule and PEMT the pre-existing microtubule.

### 2.5 Plant materials and growth conditions

The *Arabidopsis thaliana* ST-YFP and YFP-TUA5 transgenic lines were described previously [5, 43].

All seeds were sterilized with 30% bleach for 15 minutes, thoroughly washed with ddH2O, and stored at 4 °C for at least 3 days. Etiolated seedlings were grown on vertical half-strength Murashige and Skoog (MS) plates without sucrose [1/2 × MS salts, 0.8% agar, 0.05% monohydrate 2-(N-Morpholino) ethanesulfonic acid (MES), pH 5.7] in darkness at 22 °C.

For drug treatment, 2.5-day-old etiolated seedlings were carefully transferred to a fresh-made medium with drug and grown vertically for 30 minutes in darkness at room temperature. The 2 *μ*M Latrunculin B medium was prepared by adding Latrunculin B stock (2.5 mM, dissolved in DMSO) into half-strength MS plates without sucrose. The mock medium was prepared by adding same volume of DMSO.

### 2.6 Live-cell imaging and analysis

All images and videos were obtained from epidermal cells of 2.5-day-old etiolated hypocotyls, 0.5 - 1.5 mm below the apical hook. Imaging was performed on a Yokogawa CSUX1 spinning-disk system as described previously [73], obtained by Metamorph (Molecular Devices, CA) and analyzed by Fiji [68].

For imaging of YFP-TUA5, the cells chosen for analysis were all positioned two to three cells below the curved cells of the apical hook, within the region indicated previously.

For imaging of ST-YFP, 120-second time-series videos with 1-second intervals were obtained. ST-YFP particles were manually selected, traced and further analyzed by Fiji [68].

## 3 Results

### 3.1 Numerical tests verify the computational FSI model

We first conducted some relatively simple numerical experiments as a way to verify our FSI model and numerical algorithm. To that end, we set the fiber density to be slightly higher than the fluid density so that fibers moved downward due to gravity, where the gravitational force was calculated by equation (11). Consequently, the flow was predominantly unidirectional (i.e., in the vertical direction), which made it easier to examine the flow field and the fiber movement. On the other hand, the essential components of our computational model that include the nonlinear interaction between the fluid and fibers and the encounter dynamics between fibers, were fully tested.

For convenience, we assumed that all the variables and parameters are dimensionless. We considered the fluid-structure interaction of multiple elastic fibers in a box of size [0, 1]×[0, 1] containing a viscous fluid. The fibers were generated from the center of the domain via a logistic growth model:

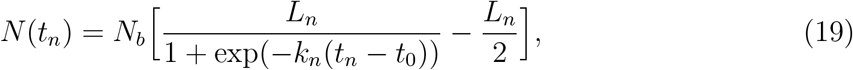

with *L*_*n*_ = 2, *t*_0_ = 0, and *n* = 1, 2, · · ·. The parameter *N*_*b*_ is the total number of fibers we intended to generate in the simulation, and *k*_*n*_ is the rate at which the fibers are generated. In the numerical implementation, the value of *N* (*t_n_*) was rounded to the nearest integer for the number of fibers at each *t_n_*. The initial length of each fiber was set to 0.05. The initial orientation of each fiber was set by a random number between 0 and 180 degrees. Each generated fiber would interact with the fluid and likely collide with other fibers. Periodic boundary conditions were applied in the vertical direction and no-slip boundary conditions were applied in the horizontal direction.

We started by running a numerical experiment in a fluid of density *ρ* = 1 and viscosity *μ* = 0.01 with a small number of fibers *N*_*b*_ = 15, and *k*_*n*_ = 0.7. We examined a typical set of results for the motion of the fibers over time (Figure 2). In particular, at *t* = 0.03125, we observed the first occurrence of the induced catastrophe. Soon after that at *t* = 0.039, a fiber was elongated due to the zippering and another was shortened due to the induced catastrophe. At the final time *t* = 0.125, a number of fibers had disappeared due to multiple zippering events.

**Figure 2:**
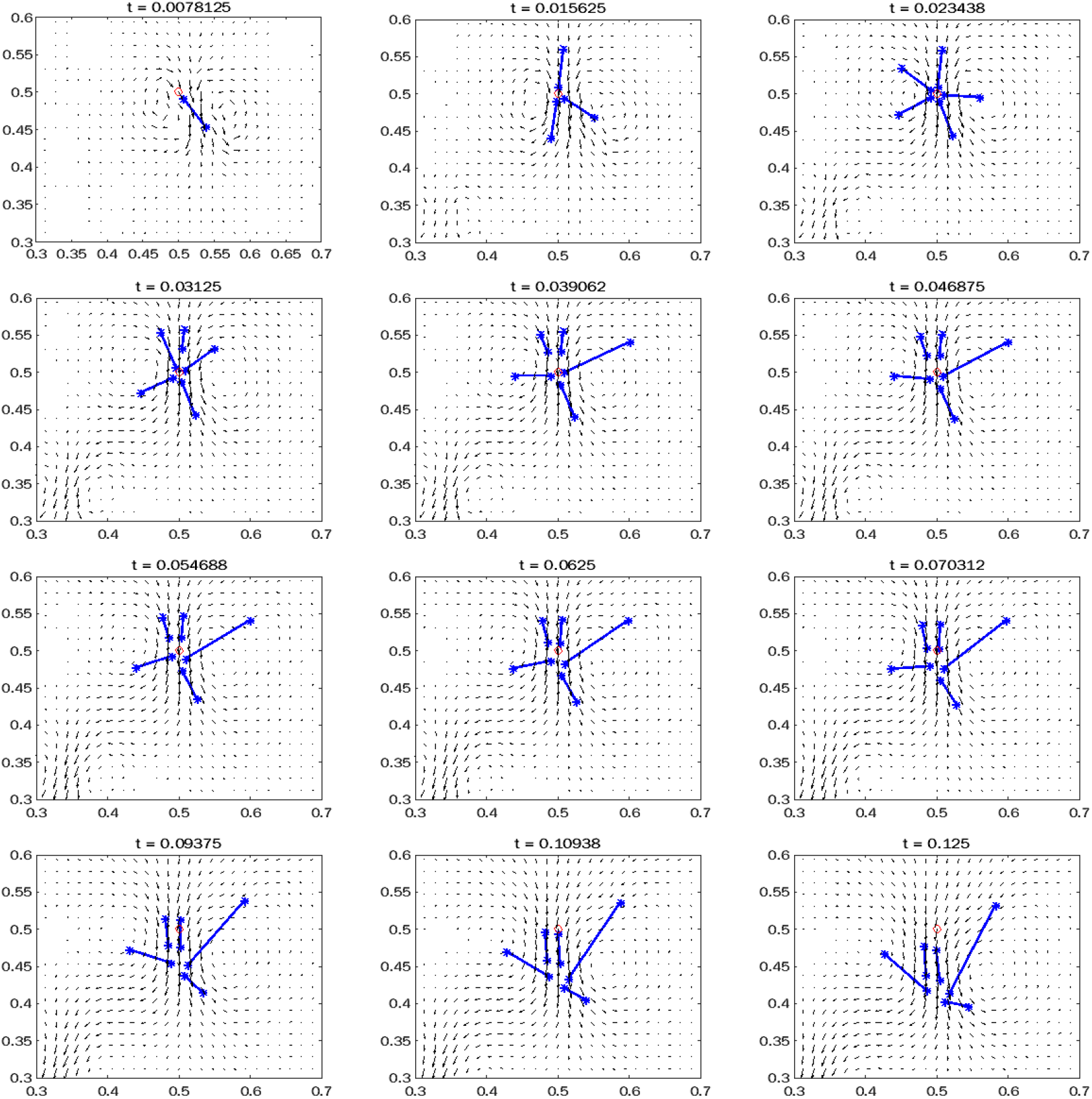
Snapshots for the FSI simulation of fibers with *N*_*b*_ = 15 at different times. In each panel, the horizontal and vertical axes represent the spatial coordinates, the red circle marks the location of fiber generation, the blue lines depict the fibers, and the small black arrows represent the fluid velocity field.

We plotted the fiber life history which depicts the variation in the length of each fiber over time (Figure 3). The horizontal lines at the bottom refer to the fibers that have disappeared due to zippering. For instance, the 10^*th*^ fiber became zippered to the 13^*th*^ fiber so that the 10^*th*^ fiber disappeared and the 13^*th*^ fiber was elongated. Some fibers experienced significant changes in their lengths as a result of either zippering or induced catastrophe, while other elastic fibers had small changes in their lengths over time, owing to their movement and deformation that resulted from the interaction with the fluid.

**Figure 3:**
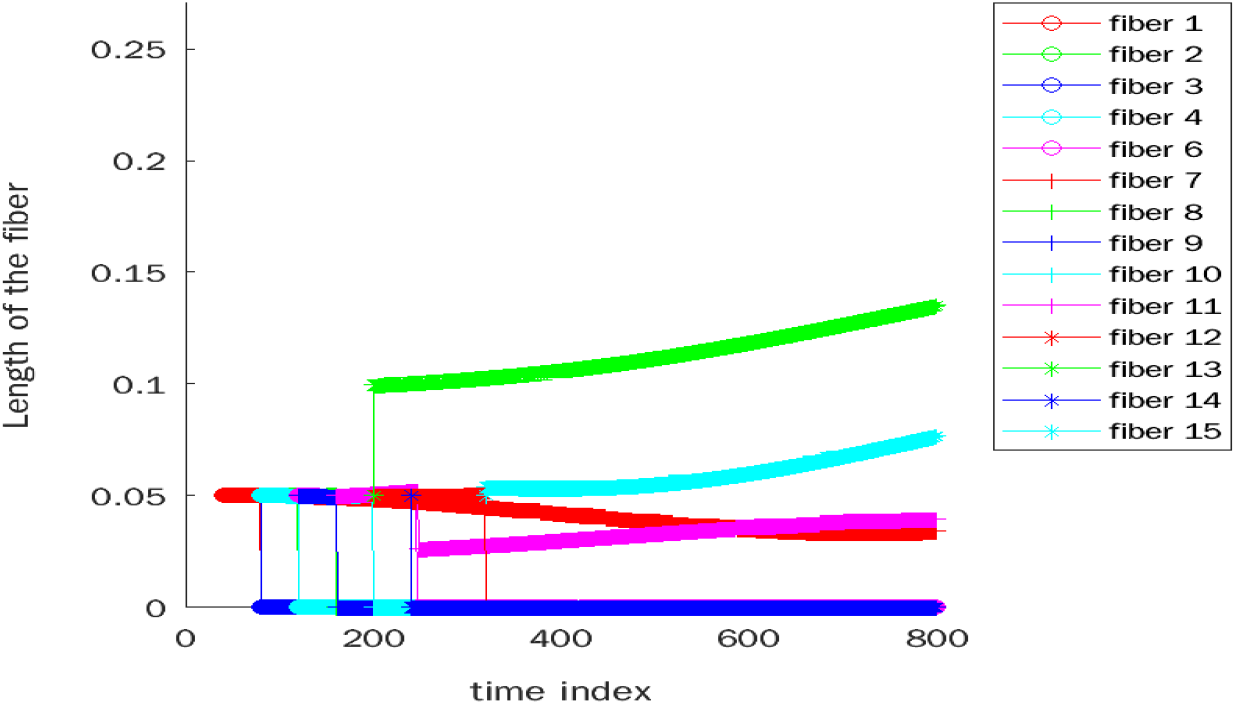
Life history of the fibers with *N*_*b*_ = 15. Zippering is seen to occur multiple times causing a significant increase in the lengths of a few fibers and the disappearance of some other fibers. Vertical lines with the fiber length dropping to zero indicate zippering.

We performed a series of numerical experiments where a few hundreds of fibers were involved in a fluid of density *ρ* = 1 and viscosity *μ* = 0.1. While it was possible to track each fiber on an individual basis in the previous example, this became challenging when a large number of fibers were present. To gain an understanding of the essential dynamics in the current numerical tests, we examined the average length and orientation of the fibers and the frequency of occurrence for zippering and induced catastrophe over time at the system level. We conducted numerical experiments with *N*_*b*_ ranging between 100 and 400. In each case, fibers were generated at the rate *k*_*n*_ = 0.1.

We studied the time evolution of the average length of the fibers with various *N*_*b*_ (Figure 4). We observed that the average fiber length experienced dynamical changes over time, due to the effects of zippering (which tends to increase the average length) and induced catastrophe (which tends to decrease the average length). A general pattern, however, was that the average length appeared to increase over time, indicating that the impact of zippering outweighed that of catastrophic collision. Such a trend is in agreement with simulation results in prior studies based on different computational models [1].

**Figure 4:**
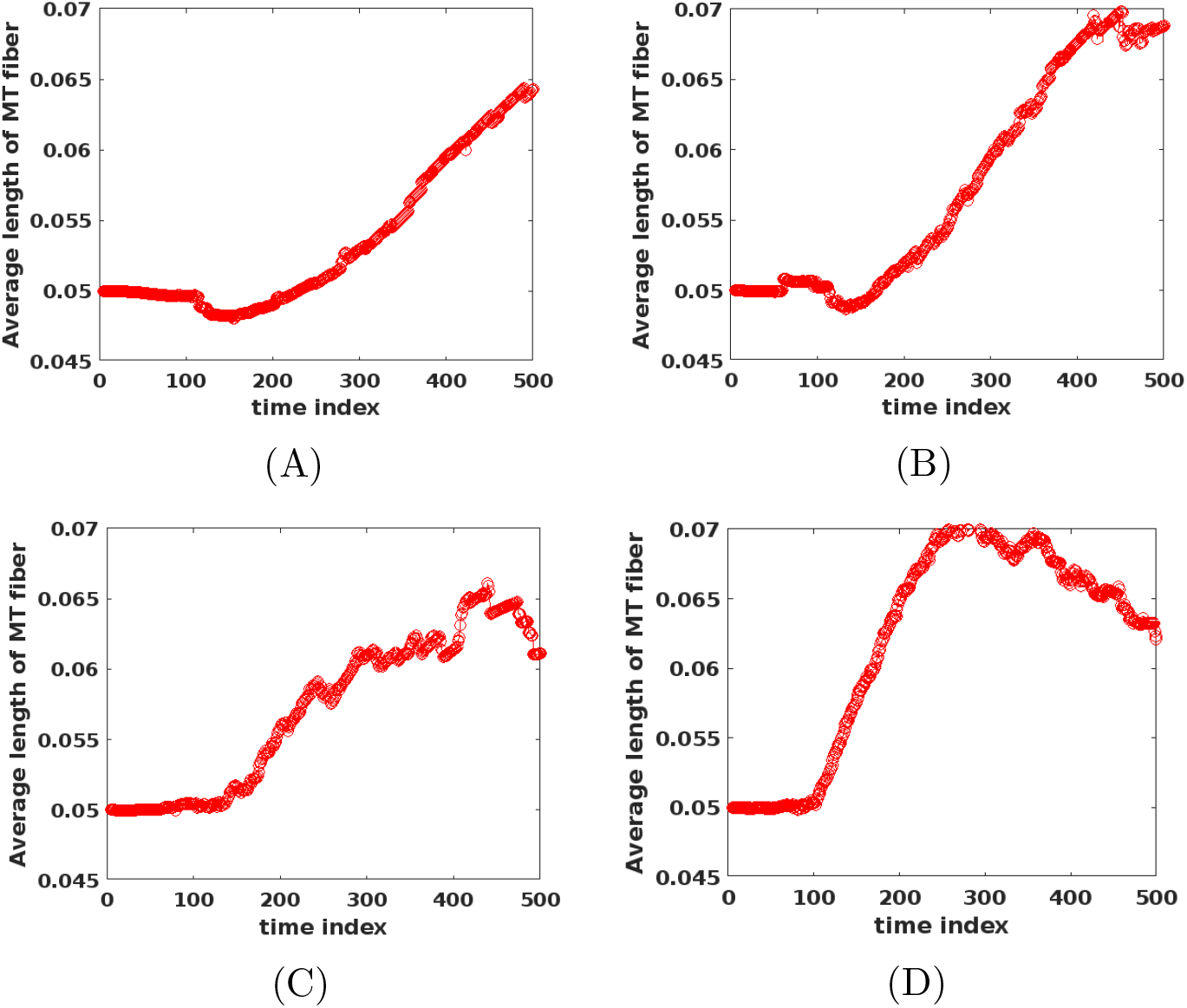
Average length of the fibers with different values of the fiber number *N_b_*. Results show that the average fiber length generally grows with time for each choice of *N*_*b*_. (A) *N*_*b*_ = 100; (B) *N*_*b*_ = 200; (C) *N*_*b*_ = 300; (D) *N*_*b*_ = 400.

We plotted the histogram, for each run, as a measure of the frequency for the relative angle between two fibers when they encounter each other (Figure 5). Angles between 0 and 40 degrees occurred much more frequently than those between 40 and 90 degrees, with angles lower than 20 degrees dominating. Such a pattern was getting slightly more pronounced as the value of *N*_*b*_ was increased. The results indicate that zippering took place much more frequently than induced catastrophe did. In particular, when *N*_*b*_ = 400, the number of occurrences for zippering was about three times that of induced catastrophe.

**Figure 5:**
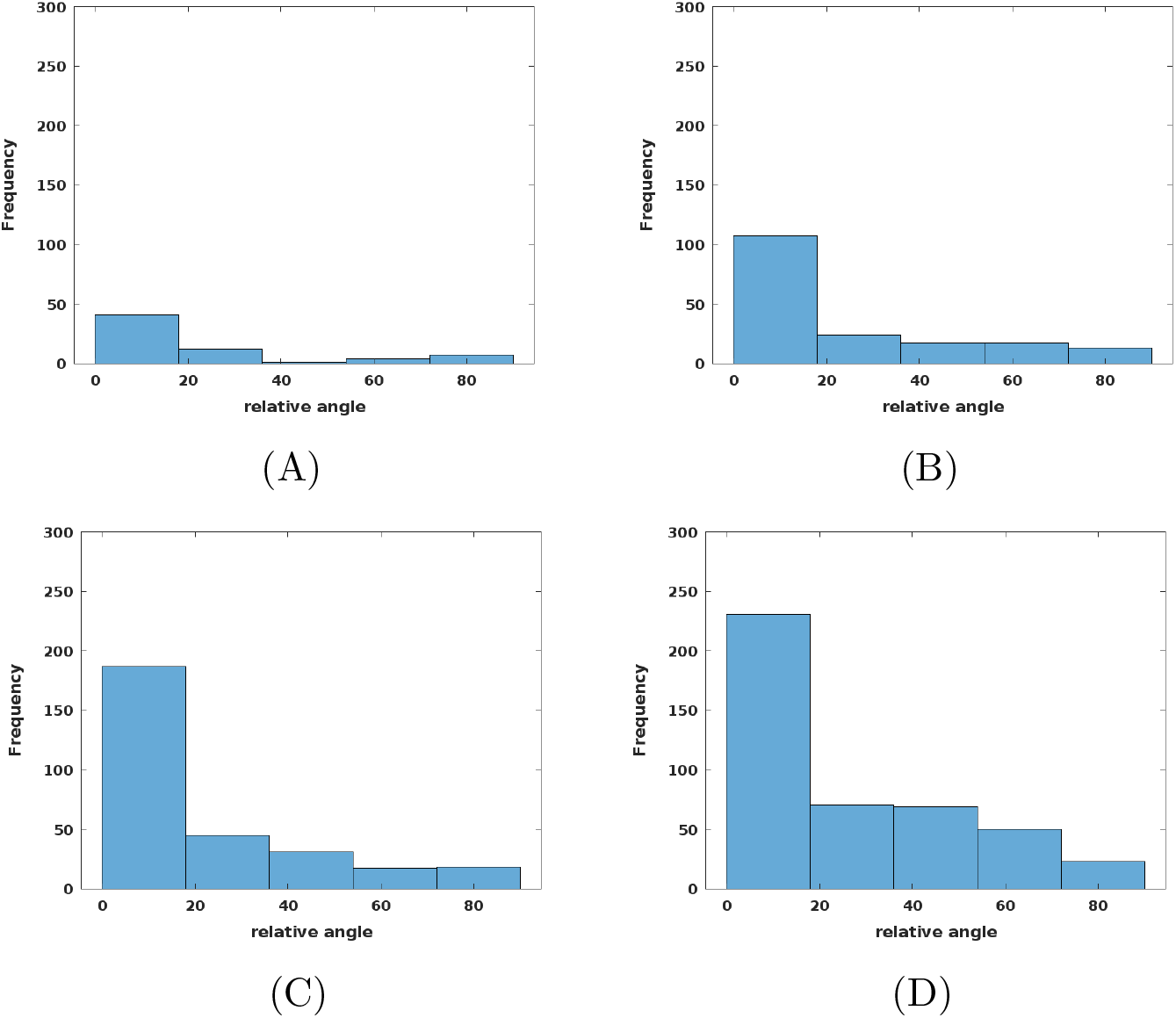
Histograms for the frequency of the relative angle formed by two microtubules when they encounter each other. Results indicate that zippering occurs more frequently than induced catastrophe. (A) *N*_*b*_ = 100; (B) *N*_*b*_ = 200; (C) *N*_*b*_ = 300; (D) *N*_*b*_ = 400.

We examined the average orientation of the fibers in reference to the flow direction. In this test, the fibers moved downward in the vertical direction due to gravity. We thus computed the angle of each fiber relative to the vertical axis in absolute value, and then took the average among all the fibers at each time. We plotted the time evolution of the average fiber orientation for different values of *N*_*b*_ (Figure 6). In most of these cases, we observed that the average angle with respect to the vertical axis decreased over time, indicating that the fibers increasingly align with the flow direction. Similar results were reported for numerical simulation of rigid fibers sedimenting in viscous flow [70], though our numerical tests here were concerned with deformable fibers.

**Figure 6:**
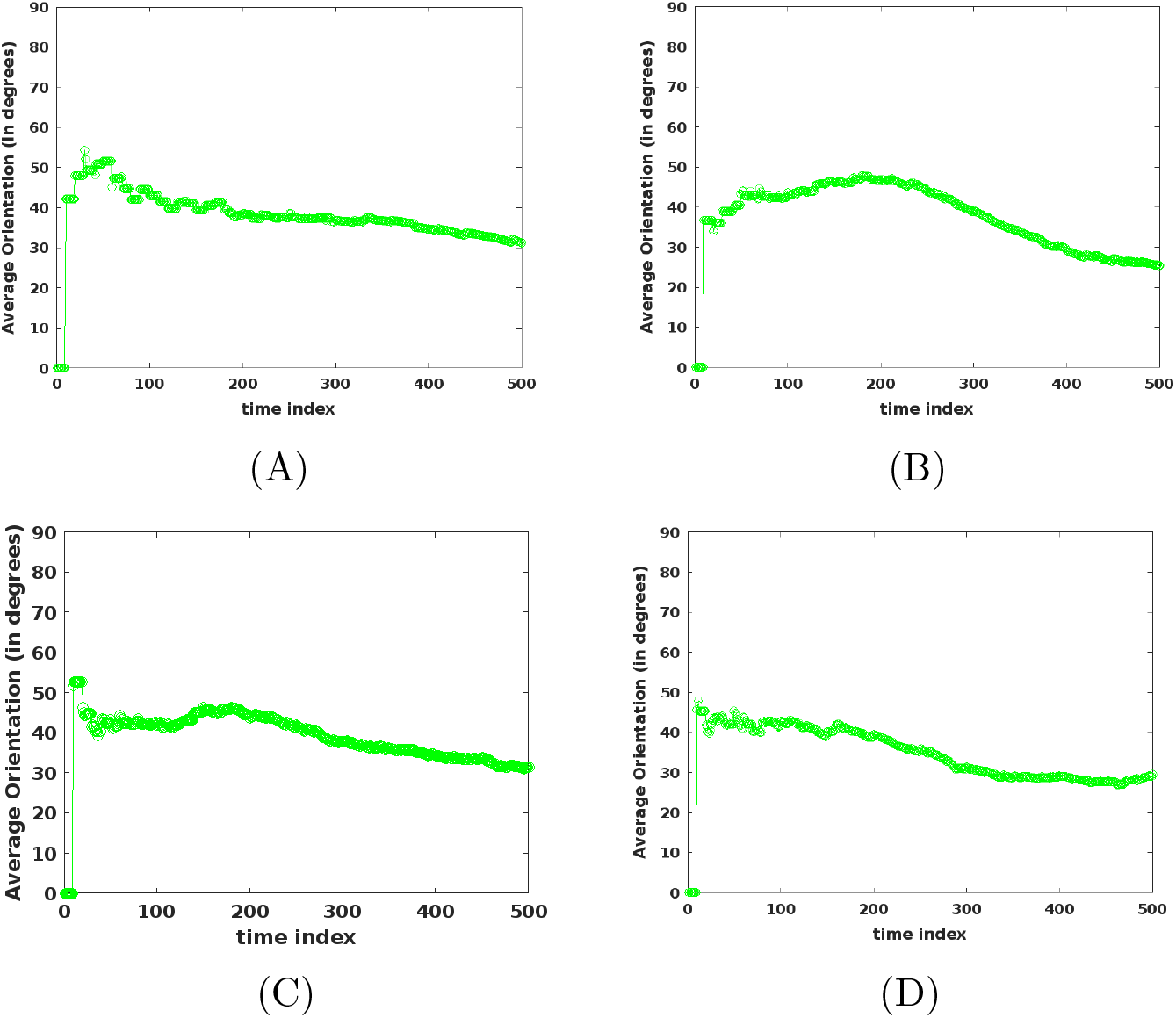
Evolution of the average orientation of the fibers in reference to the flow direction. Results show that the average angle with respect to the vertical axis tends to decrease over time. (A) *N*_*b*_ = 100; (B) *N*_*b*_ = 200; (C) *N*_*b*_ = 300; (D) *N*_*b*_ = 400.

### 3.2 Simulation shows that microtubules are primarily oriented in the direction transverse to the cell elongation axis

After we thoroughly tested our computational methods for the fiber dynamics, we conducted FSI simulation for the microtubules in a live plant cell. Our simulation setting was based on physically and biologically meaningful parameters and forces.

We set the computational domain as a rectangle of 1 mm × 0.2 mm that represented the cross section of an elongated plant cell. The horizontal direction represented the axis of cell elongation. The fluid inside the cell (i.e., cytoplasm) was treated as water, with density *ρ* = 1000 kg/m^3^ and viscosity *μ* = 0.001 Pa·s. Each microtubule was represented as a fiber with an initial length of 5 *μ*m. We assumed that the density of the microtubules was the same as that of water so that the microtubules were suspending in the fluid. The flow field was initialized with a constant velocity in the horizontal direction. The initial flow speed was set as 2.1 *μ*m/s, based on our experimental measurement on the average speed of the cytoplasmic flow (Section 3.3). No-slip conditions were applied at all the four boundary edges of the domain.

In addition to the spring force and bending force presented in equation (12), microtubules in a live plant cell are subject to a tensile stress, ***F***_*tens*_, caused by turgor pressure. The maximal tensile force acts in the direction perpendicular to the axis of cell elongation [61]. It was found that 1–2 mN is within the physiologically relevant range of the tensile stress [60]. In our simulation, we considered the effect of the maximal tensile stress only. We set ***F***_*add*_ = ***F***_*tens*_ in equation (12), where the force ***F***_*tens*_ acted in the vertical direction. We started the simulation with a force magnitude of *F*_*tens*_ = 1 mN.

We used the logistic equation (19) to model the initiation of microtubules (through a nucleation process) in the plant cell. For *N*_*b*_ = 100, we plotted the simulation result for the frequency of the encountering angles (Figure 7A). We observed that angles between 0° and 40° still dominated, a pattern similar to that in Figure 5, indicating a more frequent occurrence of zippering. On the other hand, angles larger than 40° took a higher percentage in comparison to that in Figure 5, indicating that induced catastrophe had a higher frequency of occurrence in a real plant cell, possibly due to the impact of the tensile force.

**Figure 7:**
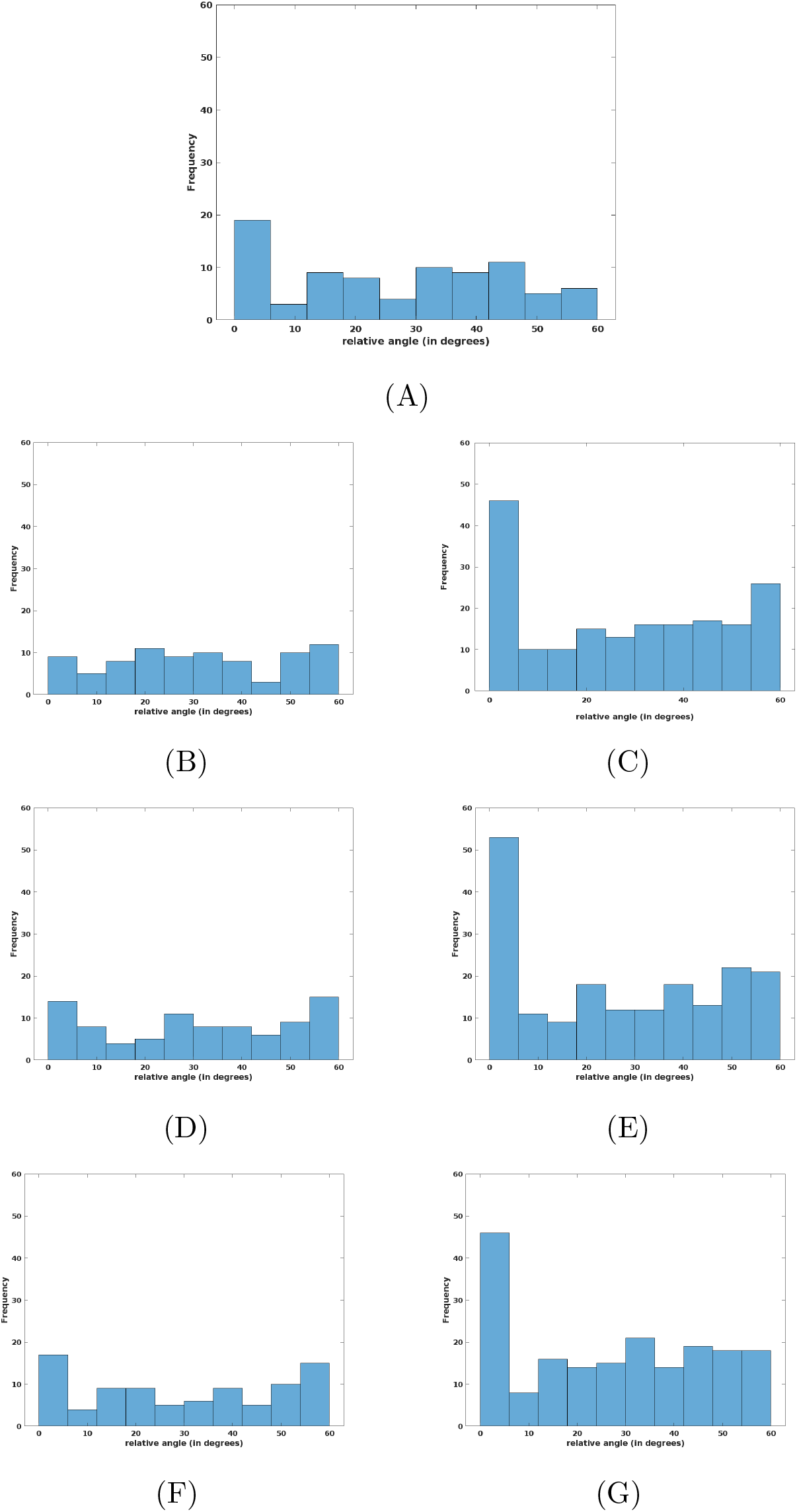
Frequency of the encountering angles for the microtubules with different values for the magnitude of the tensile force (*F*_*tens*_) and the number of microtubules (*N*_*b*_). Results indicate that zippering occurs more frequently than induced catastrophe. (A) *F*_*tens*_ = 1 *mN*, *N*_*b*_ = 100; (B) *F*_*tens*_ = 2 *mN*, *N*_*b*_ = 100; (C) *F*_*tens*_ = 2 *mN*, *N*_*b*_ = 200; (D) *F*_*tens*_ = 5 *mN*, *N*_*b*_ = 100; (E) *F*_*tens*_ = 5 *mN*, *N*_*b*_ = 200; (F) *F*_*tens*_ = 10 *mN*, *N*_*b*_ = 100; (G) *F*_*tens*_ = 10 *mN*, *N*_*b*_ = 200.

Meanwhile, we plotted the frequency of the MT orientation, measured in the angle formed by each microtubule with respect to the horizontal axis (Figure 8A). As can be clearly seen, angles between 80° and 90° took a dominant role, showing that the vast majority of the microtubules aligned with the vertical axis that was perpendicular to the direction of cell elongation. In our previous tests where the flow was mainly driven by gravity, we found that the average fiber orientation decreases to 40° or below (Figure 6), an indication that most fibers tend to align themselves in the vertical direction (i.e., the flow direction). For our current simulation of MT dynamics, with microtubules suspending and floating in cytoplasm, the flow field was more complex than being unidirectional. In fact, the fluid flow involved significant movement in both the horizontal and vertical directions (see Section 3.3). The results in Figure 8A, however, showed that the microtubules were still primarily oriented along the vertical direction, which corresponded to the direction of the maximum tensile stress [61, 66, 67].

**Figure 8:**
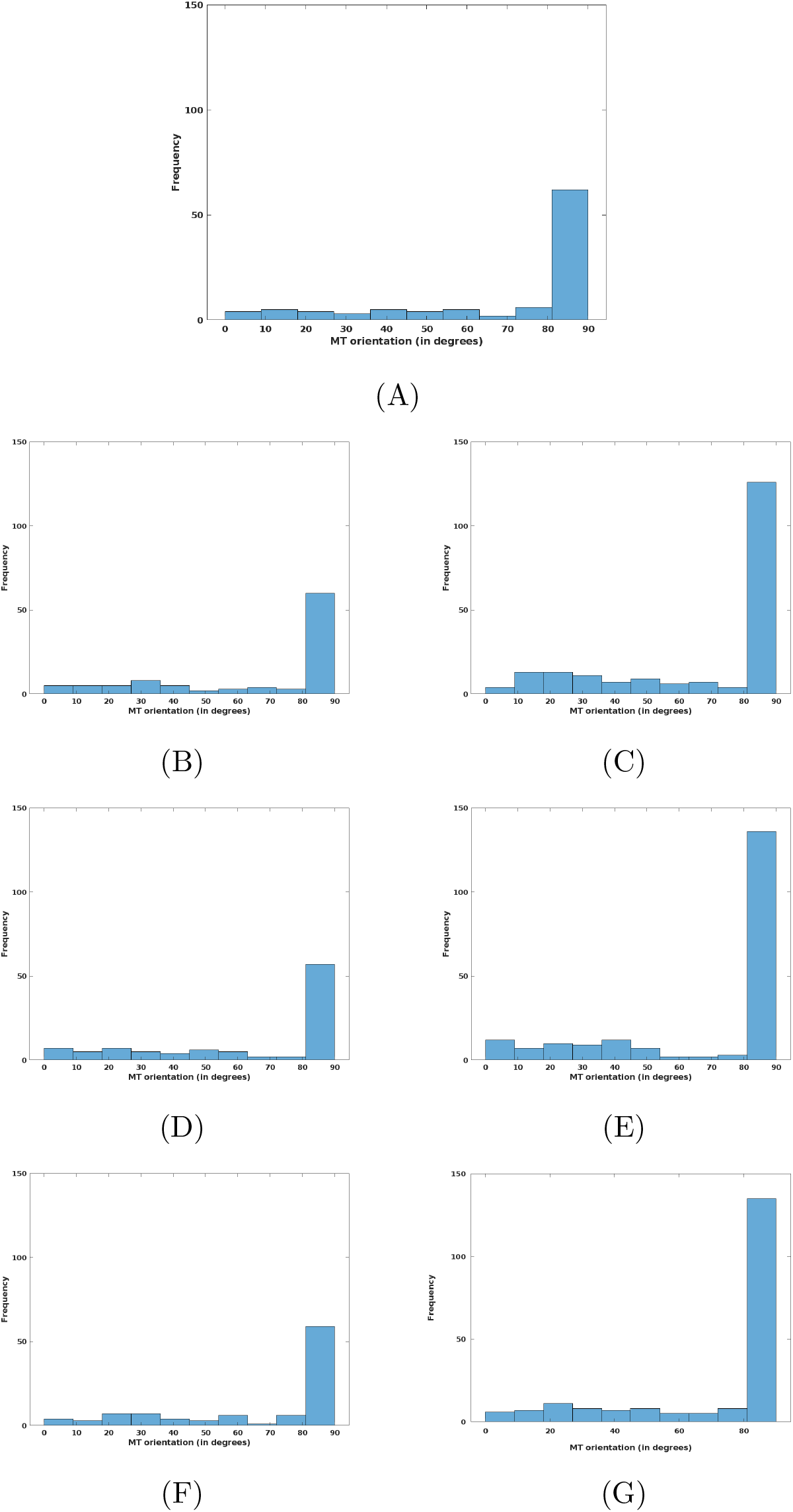
Frequency of the orientation angles for the microtubules with different values for the magnitude of the tensile force (*F*_*tens*_) and the number of microtubules (*N*_*b*_). Results show that the microtubules are primarily oriented along the vertical direction, corresponding to the direction of the maximum tensile stress. (A) *F*_*tens*_ = 1 *mN*, *N*_*b*_ = 100; (B) *F*_*tens*_ = 2 *mN*, *N*_*b*_ = 100; (C) *F*_*tens*_ = 2 *mN*, *N*_*b*_ = 200; (D) *F*_*tens*_ = 5 *mN*, *N*_*b*_ = 100; (E) *F*_*tens*_ = 5 *mN*, *N*_*b*_ = 200; (F) *F*_*tens*_ = 10 *mN*, *N*_*b*_ = 100; (G) *F*_*tens*_ = 10 *mN*, *N*_*b*_ = 200.

Next, we conducted the simulation with different numbers of microtubules and different magnitudes of the tensile force acting on each microtubule. We plotted the frequency for the encountering angles with the magnitude of the tensile force *F*_*tens*_ = 2 mN, 5 mN, and 10 mN (Figure 7B-G). For each case, we examined *N*_*b*_ = 100 and *N*_*b*_ = 200. The results are similar to those in Figure 7A. We also plotted the frequency for the MT orientation angles under the same settings (Figure 8B-G). Again, we observed a similar patten to that in Figure 8A; i.e., the vast majority of microtubules aligned themselves in the vertical direction. This pattern was getting more pronounced as either the force magnitude or the number of microtubules was increased. Nevertheless, even with the presence of a tensile force (e.g., 10 mN) that was significantly larger than normal, a portion of microtubules always exhibited an orientation angle ranging between 0° and 60°, indicating the impact of fluid flow in directions different from that of the maximum tensile stress. We emphasize, however, that the effects of the tensile force and fluid dynamics are not separated. As the tensile stress is acting on microtubules, the force is transmitted from microtubules to the fluid through the FSI, which then impacts the motion of the fluid. Subsequently, the fluid flow impacts the movement of microtubules because of the no-slip condition, providing a feedback to the MT dynamics. Although the alignment of microtubules with the maximal tensile stress direction has been observed in many experimental studies, how this is achieved still remains largely unknown [61, 67]. Our simulation results suggest that the fluid (cytoplasm) may serve as a medium to transmit the force and to regulate the MT alignment. The overall pattern of the MT assembly and orientation is thus shaped by the combined effects of the fluid flow, the tensile force, and the interaction between microtubules and cytoplasm.

### 3.3 Cytoplasmic flow impacts the microtubule orientation in live cells

To validate our computational model for the MT dynamics, we conducted laboratory experiments in the *Arabidopsis thaliana* etiolated seedlings. The focus of these experiments was to monitor the cytoplasmic flow and to examine how it would impact the MT orientation. The results would provide experimental measurements to compare with our computational predictions for the interaction between microtubules and cytoplasm. Meanwhile, since the tensile stress is always present in a real plant cell, the experimental findings would reflect the impact of the tensile force on the MT dynamics.

We analyzed the movement of the ST-YFP fusion proteins at the cortical cytoplasm, which represents the dynamics of Golgi and Golgi stacks, as a proxy for the cytoplasmic flow [5]. We found that the Golgi stacks were highly motile with a wide range of speeds (Figure 9A). The maximum speed could exceed 7 *μ*m/sec. The speed of Golgi could reduce from the maximum speed to a complete stop in a few seconds as Golgi shifts between saltatory and continuous movement rapidly (Figure 9B and Movie S1).

**Figure 9:**
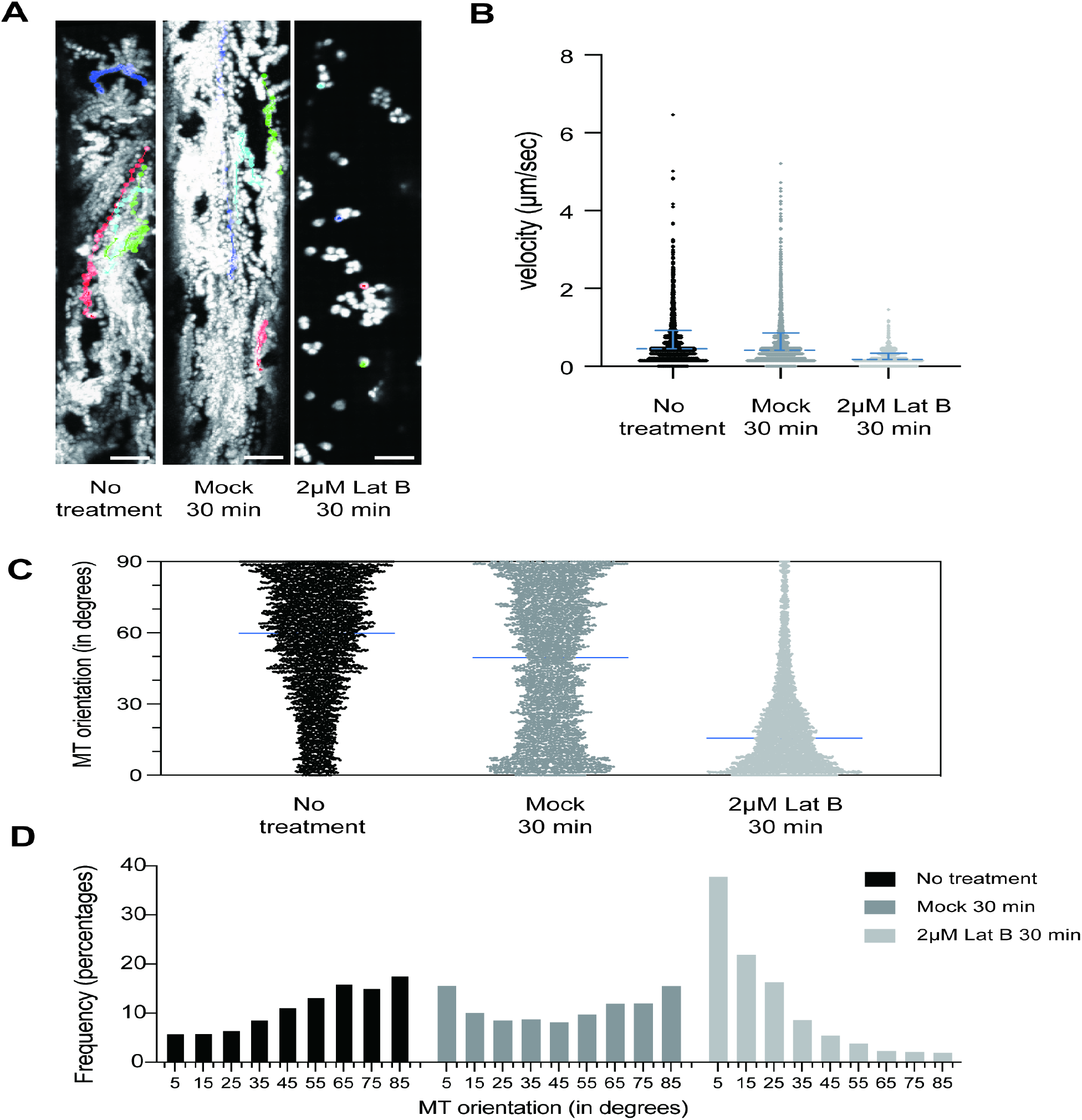
Impact of cytoplasmic streaming on the mobility of Golgi and the orientation of microtubules. (**A**) Representative trajectories of Golgi under no treatment, mock treatment, and 2 *μ*M Latrunculin B (Lat B) for 30 min. The Z-stacks were obtained from 2 min videos. Bar = 5*μ*m. (**B**) Analysis of the velocity of Golgi under no treatment, mock treatment, and 2 *μ*M Lat B for 30 min. Data were collected from 100 tracks with at least 20 cells from 7 seedlings for each condition. Bar represents the mean velocity with SD. (**C**) Analysis of the MT orientation under no treatment, mock treatment, and 2 *μ*M Lat B for 30 min. About 4000 reads of microtubules were collected from 40 cells from 7 seedlings for each condition. Bar represents the median angle of the MT orientation. (**D**) Histograms of the MT orientation angles under no treatment, mock treatment, and 2 *μ*M Lat B for 30 min.

We analyzed the MT arrangements under the same growth condition using YFP-TUA5 transgenic lines, which marked an isoform of *α*-tubulin [43]. We found that 48.72% of the microtubules were oriented between 60° and 90° and the median was 59.04° (Figure 9C and 9D), where a 90° microtubule was regarded as perpendicular to the cell elongation.

To manipulate the cytoplasmic flow of the plant cell, we applied Lat B, which is commonly used to pause the cytoplasmic streaming [6]. 2.5-day-old etiolated seedlings of ST-YFP lines were treated with 2 *μ*M Lat B for 30 minutes. We recorded the dynamics of Golgi stacks after the Lat B treatment and found it significantly reduced the dynamics of Golgi stacks, particularly the saltatory movement (Figure 9A and 9B, Movie S2 and S3). We examined the microtubule orientation under Lat B treatment using YFP-TUA5 lines. The median orientation angle of microtubules was 15.52° under the Lat B treatment, due to the fact that most microtubules (75.89%) were oriented between 0° and 30°. This is significantly different from microtubules under the mock treatment where the median orientation angle was 48.78° and where 39.37% of the microtubules were oriented between 60° and 90° (Figure 9C and 9D). Thus, the in vivo observation indicates that the cytoplasmic flow plays a significant role in determining the orientation of MT, which is consistent with the computational prediction.

### 3.4 Integration of computational modeling and experimental testing helps to refine the model

To integrate the computational modeling and experimental testing, we compared datasets generated by the numerical simulations and biological experiments. The simulation result in the base setting (*F*_*tens*_ = 1 *mN*, *N*_*b*_ = 100) showed that about 64% of the microtubules were oriented between 60° and 90° (Figure 8A), higher than what was obtained from the experimental measurement (48.72%, Figure 9D). Moreover, there were many more microtubules in the experiment than in the computation that were aligned between 0° and 60°. One possible reason for this discrepancy was that we utilized a simplistic treatment of the tensile stress in our computational model. We only considered the maximal tensile force, which acted in the vertical direction (perpendicular to the axis of cell elongation). In a real plant cell, the tensile stress is also present in other directions, which could impact the orientation of the microtubules.

To better represent the effects of the tensile stress, we kept the magnitude of the maximal tensile force at 1 mN and added a secondary tensile force into our computational model, which acted in the horizontal direction with a magnitude of 0.4 mN. We then ran the simulation again with *N*_*b*_ = 100, and presented the result in Figure 10. We observed that about 49% of the microtubules were oriented between 60° and 90°, indicating a very good agreement with the percentage observed from the experiment.

**Figure 10:**
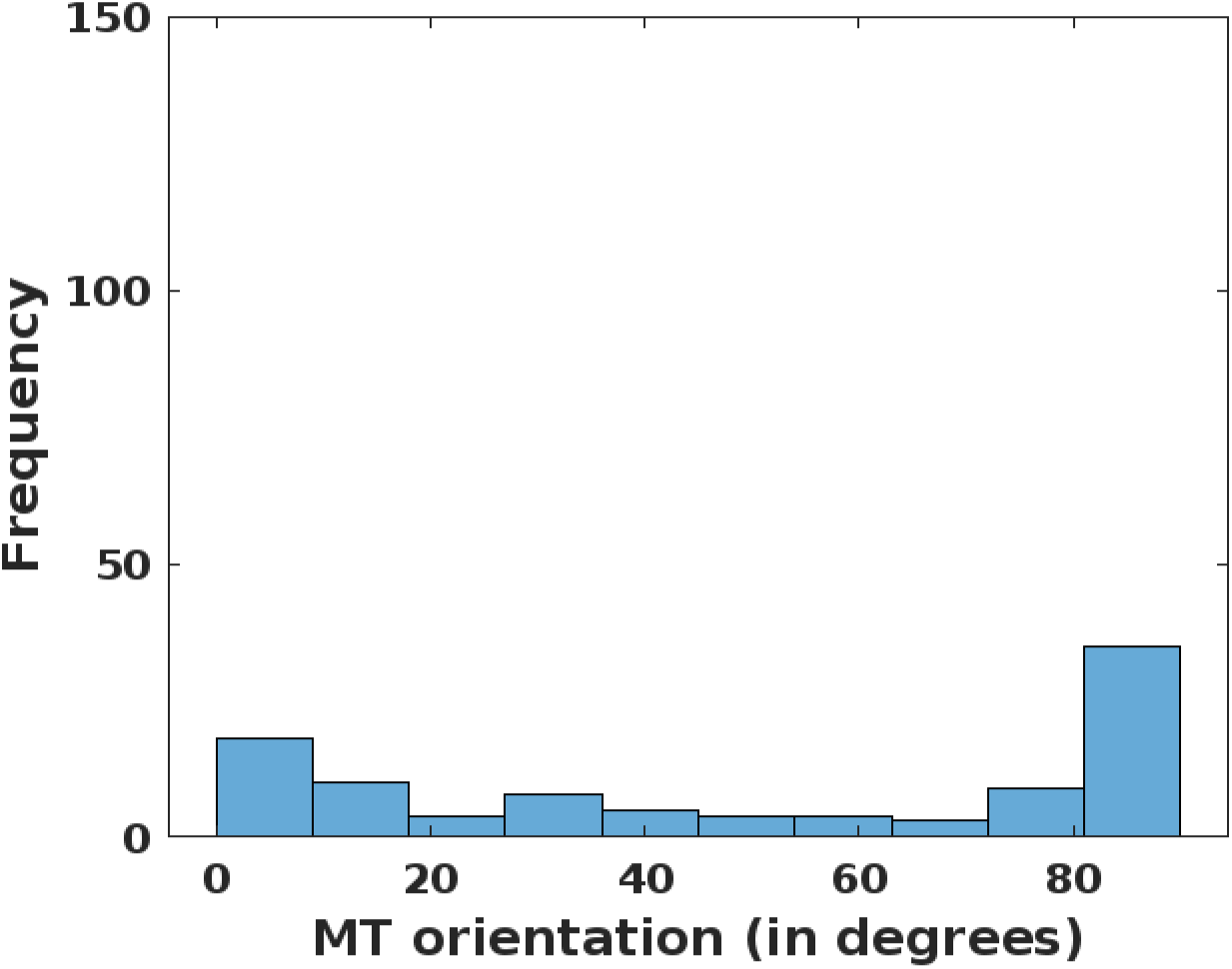
Frequency of the orientation angles for the microtubules with *N*_*b*_ = 100. The maximal tensile force acts in the vertical direction with a magnitude of 1 mN, and the secondary tensile force acts in the horizontal direction with a magnitude of 0.4 mN.

## 4 Discussion

We have developed a mathematical and computational framework to investigate the interaction between microtubules and cytoplasm within plant cells, in order to test our hypothesis that such an interaction plays a significant role in regulating the spatial organization of microtubules. We have utilized the theory, modeling techniques and numerical algorithms from fluid-structure interaction in this study. In particular, the immersed boundary method is employed to conduct the FSI simulation for microtubules. Details of the MT encounter dynamics are also incorporated into our computational study. Results are presented for multiple fibers interacting with viscous fluid flow as numerical tests for our model and methods, and for the simulation of MT dynamics in a real plant cell. The MT simulation results are also compared to the measurement from laboratory experiments.

Our numerical simulation has explored a range of parameter settings that include the number of microtubules and the magnitude of the tensile force. We have paid special attention to the orientation of microtubules and the frequency of zippering and induced catastrophe in the model output. The numerical tests in Section 3.1 and the MT simulation in Section 3.2 share several common results, especially in the frequency of occurrence for zippering and induced catastrophe. It is found that zippering occurs more frequently than induced catastrophe for both scenarios, in spite of the different settings and different values of the physical parameters such as the density and viscosity. However, the simulation results for the fiber orientations in these two scenarios are distinct from each other: the fibers in the numerical tests tend to align in the direction of the gravity-driven flow, whereas the microtubules suspending and moving in the more complex cytoplasmic flow are primarily oriented in the direction perpendicular to the axis of cell elongation. This inherent difference is largely due to the presence of the tensile stress acting on the microtubules in a real plant cell. Nevertheless, the impacts of the tensile stress (as well as other forces acting on microtubules) and fluid dynamics are coupled together through the fluid-structure interaction, which seems to form a stress-microtubule-cytoplasm nexus. Consequently, the overall pattern of the MT organization and orientation appears to be heavily influenced by the interconnected microtubule-cytoplasm system and the communication of forces and velocities between each other. We have simulated different numbers of microtubules and different magnitudes of the tensile force within a biologically feasible regime, and all the computational findings exhibit a consistent pattern. We have also found that, with additional calibration of the tensile stress computation, our simulation outcome could lead to a good agreement, both qualitatively and quantitatively, with the experimental measurement.

We have focused on the impact of fluid dynamics on MT assembly and orientation in the present paper. As such, we have not considered other intracellular components such as motor proteins, enzymes, chromosomes, organelles and other endosomes, which also interact with microtubules and may contribute to their spatial organization as well. Meanwhile, our model is primarily deterministic and does not take into account some of the stochastic behaviors, such as treadmilling, of microtubules. These factors might have contributed to the quantitative differences between our computational and experimental results on the MT orientation. Nevertheless, our simulation results suggest that an FSI study incorporating realistic forces and encounter dynamics would be sufficient to generate some of the essential features of MT dynamics within a plant cell. Our modeling framework can be extended to include those additional intracellular factors and properties associated with microtubules, but we expect that the qualitative outcome would remain the same; i.e., fluid dynamics play an important role in shaping the spatial organization of microtubules.

Cytoplasmic streaming generates active cytoplasmic movement to execute many functions such as cell locomotion for nutrient uptake in unicellular organism amoeba, chloroplast movement for light response in plant cells, information exchange between intracellular organelles in mammalian and plant cells and pattern formation in *Caenorhabditis elegans* [26,28]. Cytoplasmic streaming is driven by cytoskeleton-dependent motor proteins – microtubule-based motor protein kinesin and actin-based motor protein myosin [23, 69]. The fluid dynamics of cytoplasmic streaming itself is presumed to impact the organization of actin and/or microtubule cytoskeleton. Yet the detail for the impact of cytoplasmic streaming on the cytoskeleton is poorly understood. Our results combined computational modeling and *in vivo* studies to reveal that microtubule-cytoplasmic interaction is important for maintaining proper microtubule patterning in *Arabidopsis* epidermal cells. Short treatment of Lat B reduced cytoplasmic flow, resulted in reduced mobility of Golgi, due to the impairment of actin cables along which Golgi tracks. Microtubules orientation under Lat B treatment had a 30-degree deviation from that of mock treatment. It is known that actin influences global cellulose synthase distribution and therefore it may impact cellulose microfibril patterning [34]. The microtubule reorientation, however, is unlikely impacted by actin’s role in cellulose synthase trafficking. The feedback regulation from cellulose microfibril patterning to the organization to microtubules takes days if not hours. To complement the pharmacological methods, future experimental studies will focus on disrupting genes that specifically drive the cytoplasmic streaming without minimal perturbation of actin cytoskeleton.

This work represents a pilot study for the computational modeling of fluid dynamics and MT organization in a plant cell. Although we have focused on a 2D setting, the modeling techniques and computational methods can be applied to the 3D space. MT simulation in 3D would allow us to incorporate more detailed encounter dynamics of microtubules, including the effect of cross-over that is not present in 2D, and more complex patterns of cellular fluid dynamics. We plan to pursue this research direction in our future efforts. In addition, the modeling and computational methodology based on fluid-structure interaction can be naturally extended to many other cellular systems where the intracellular fluid interacts with various microstructures and where such interaction may be important in shaping the cellular and molecular dynamics.

