## Supplemental video for "Modeling Microtubule-Cytoplasm Interaction in Plant Cells"

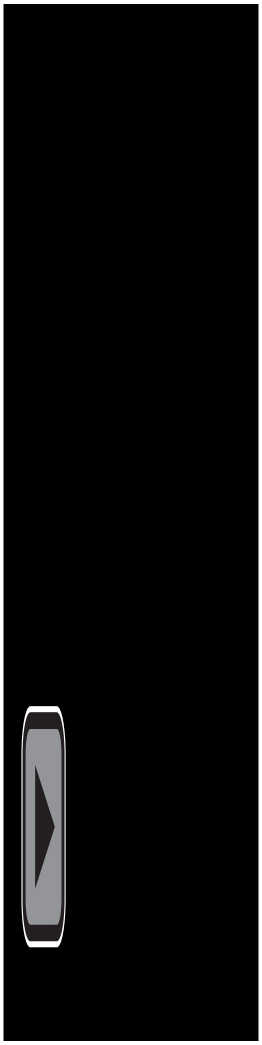

Movie S1

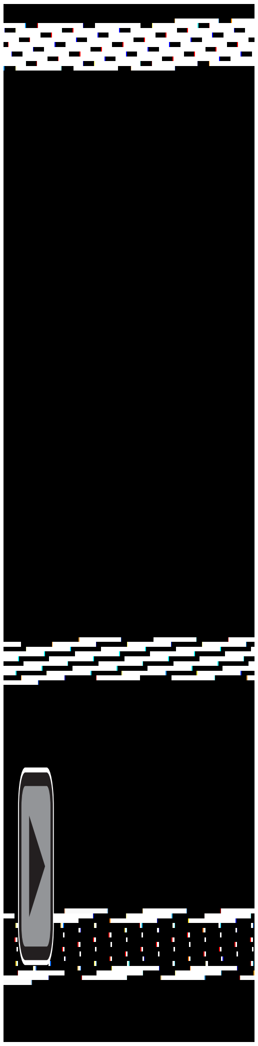

Movie S2

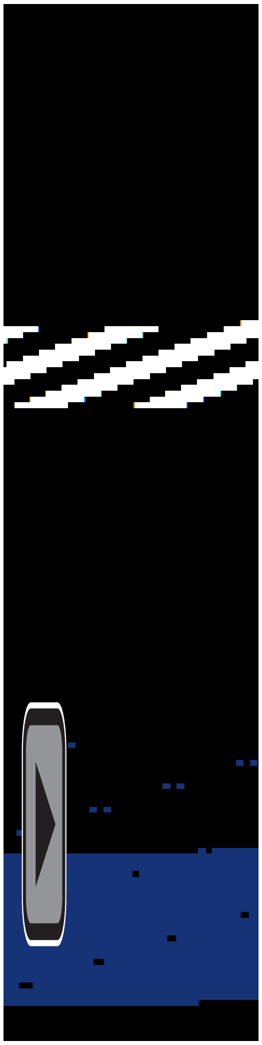

Movie S3

**Movie S1:**
Dynamics of the ST-YFP particles at the cortical cytoplasm of the Arabidopsis etiolated hypocotyl cells. The 2.5-d-old etiolated Arabidopsis seedlings expressing ST-YFP were imaged by confocal microscopy. Time series is 1-s intervals. Total time = 120 s. Frame rate = 20 fps. (Bar = 5 μm.)

**Movie S2:**
Dynamics of the ST-YFP particles at the cortical cytoplasm of the Arabidopsis etiolated hypocotyl cells. The 2.5-d-old etiolated Arabidopsis seedlings expressing ST-YFP were imaged by confocal microscopy after 30min DMSO treatment. Time series is 1-s intervals. Total time = 120 s. Frame rate = 20 fps. (Bar = 5 μm.)

**Movie S3:**
Dynamics of the ST-YFP particles at the cortical cytoplasm of the Arabidopsis etiolated hypocotyl cells. The 2.5-d-old etiolated Arabidopsis seedlings expressing ST-YFP were imaged by confocal microscopy after 30min 2μM Lat B treatment. Time series is 1-s intervals. Total time = 120 s. Frame rate = 20 fps. (Bar = 5 μm.)
